# Alignment, Segmentation and Neighborhood Analysis in Cyclic Immunohistochemistry Data Using CASSATT

**DOI:** 10.1101/2022.08.29.504975

**Authors:** Asa A. Brockman, Rohit Khurana, Todd Bartkowiak, Portia L. Thomas, Shamilene Sivagnanam, Courtney B Betts, Lisa M. Coussens, Christine M. Lovly, Jonathan M. Irish, Rebecca A. Ihrie

## Abstract

Cyclic immunohistochemistry (cycIHC) uses sequential rounds of colorimetric immunostaining and imaging for quantitative mapping of location and number of cells of interest. In addition, cycIHC benefits from the speed and simplicity of brightfield microscopy for data collection, making the collection of entire tissue sections and slides possible at a trivial cost compared to other high dimensional imaging modalities. However, large cycIHC datasets (greater than 50 GB) currently require an expert data scientist to concatenate separate open-source tools for each step of image pre-processing, registration, and segmentation, or the use of proprietary software. Here, we present a unified and user-friendly pipeline for processing, aligning, and analyzing cycIHC data - Cyclic Analysis of Single-Cell Subsets and Tissue Territories (CASSATT). CASSATT registers scanned slide images across all rounds of staining, segments individual nuclei, and measures marker expression on each detected cell. Beyond straightforward single cell data analysis outputs, CASSATT explores the spatial relationships between cell populations. By calculating the logodds of interaction frequencies between cell populations within tissues and tissue regions, this pipeline helps users identify populations of cells that interact - or do not interact - at frequencies that are greater than those occurring by chance. It also identifies specific neighborhoods of cells based on the assortment of neighboring cell types that surround each cell in the sample. The presence and location of these neighborhoods can be compared across slides or within distinct regions within a tissue. CASSATT was first tested using a newly generated cycIHC dataset consisting of six GBM tissue sections processed through eight cycles of AEC based IHC staining. Further validation was completed on a previously published lung cancer tissue microarray dataset consisting of 107 cores processed through eighteen cycles of staining and imaging. CASSATT is a fully open-source workflow tool developed to process cycIHC data and will allow greater utilization of this powerful staining technique.

## Introduction

As the use of transcriptomics and multi-omic approaches to describe cell identity becomes increasingly prevalent, an accompanying problem arises of how to identify cells of interest using complementary, low-dimensional approaches that can be readily applied to large sets of patient samples. While highly multiplexed imaging experiments using platforms such as imaging mass cytometry or digital spatial profiling are data-rich, they require advanced experimental approaches, specialized imaging equipment, and additional time when compared to standard immunostaining approaches used in the pathology laboratory. Additionally, after cells of interest are identified in high-dimensional data, it is often desirable to confirm the presence of these cells in a second, prospective sample set, typically one representing a larger number of patients or conditions. In these cases, cyclic immunohistochemistry, which uses sequential rounds of colorimetric immunostaining and imaging, can be a useful approach for quantitative mapping of location and number of cells of interest. In addition, cyclic immunohistochemistry benefits from the speed and simplicity of brightfield microscopy for data collection, making the collection of entire tissue sections a trivial cost compared to other high dimensional imaging modalities. However, such large datasets currently require an expert data scientist to bring together separate tools for each step of image pre-processing, registration, and segmentation, or expensive proprietary software such as HALO or Visiopharm (1,2). An area of particular and growing interest in the field of multiplexed imaging of solid tissues and tumor samples is the spatial relationships between cells that are expert- or machine-identified as different populations within the tissue. Example questions of interest include whether cells of one population are found to have neighbor cells of another population in greater proportion than expected by chance, given their frequency within the sample (3). An accompanying question would be whether there are assortments of neighbor cells that consistently arise within a sample - that is, multiple populations that tend to co-locate (4). Here, we present a unified and user-friendly pipeline for processing, aligning, and analyzing cycIHC data in these fashions - Cyclic Analysis of Single-Cell Subsets and Tissue Territories (CASSATT).

The objective of this analysis pipeline is to streamline, automate, and increase throughput of cyclic immunohistochemistry data analysis using open-source python packages. Here, we apply the pipeline to a test dataset of six glioblastoma tissue sections cyclically stained for eight biomarkers for a total of forty-eight scanned slide images. In brief, the steps accomplished are: image import, tissue-level registration between imaging rounds, cell-level registration, image segmentation, and modular options for neighborhood analysis (**Figure 1**). This approach provides an efficient and scalable workflow to answer many questions that are commonly encountered in discovery research in tissue specimens, including the overall abundance of cells of interest within a tissue section as well as whether specific, user-defined groups of cells are in close proximity - i.e., whether they may interact at a higher-than-random rate. CASSATT gathers together and streamlines all the major steps necessary to produce single cell expression information from cyclic immunohistochemistry datasets and does so in an open-source environment. In addition, it systematically analyzes the spatial relationships between cell populations and via unsupervised algorithms, identifies clusters of cell niches.

**Figure 1.**
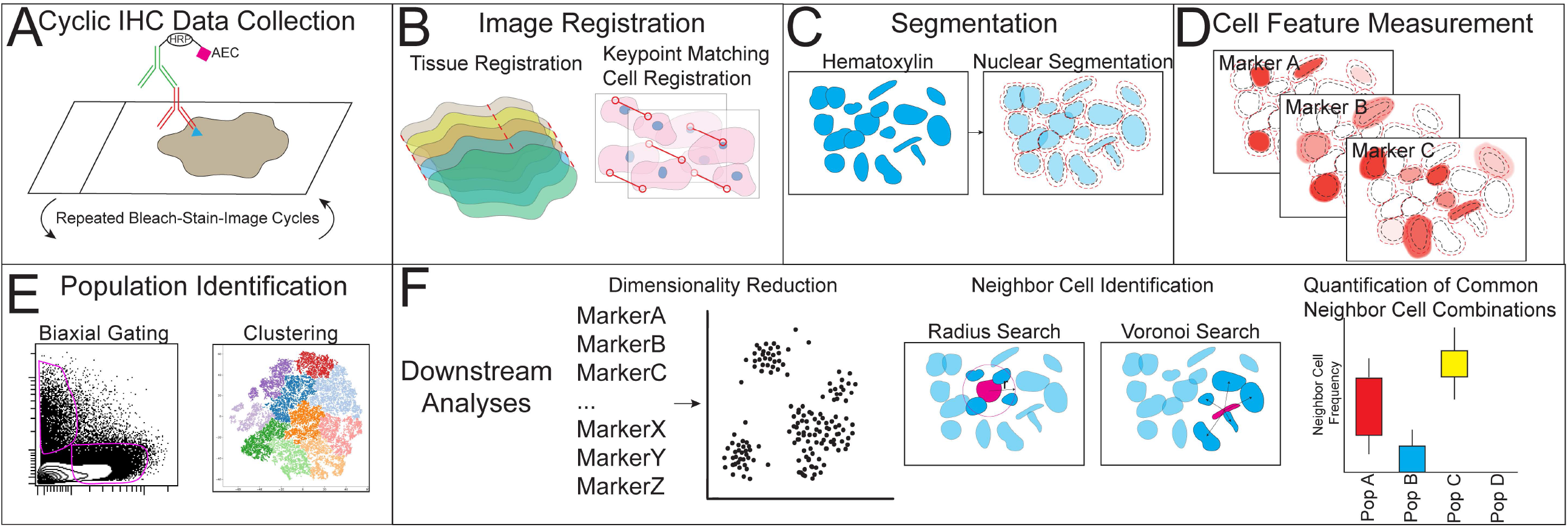
Pipeline Overview (A) Cyclic IHC is performed by repeated bleach, stain, image cycles on a single piece of tissue utilizing the amino ethyl carbazol (AEC) chromogen and a whole slide scanning microscope to acquire images. (B) Images are registered at both tissue and cell level to allow for single-cell quantification of features. (C) Nuclei are segmented using the unmixed hematoxylin stain. (D) Cell features are measured from each of the staining cycles. (E) Populations of cells are identified using either traditional biaxial gating or automated clustering techniques (F) Downstream analysis is performed including outputs of population frequency, dimensionality reduction, neighbor cell identification, and quantification of neighbor cell frequencies.

## Methods

Cyclic IHC was performed on 5 micron FFPE tissue sections according to a modified protocol based on previous studies (5-7). Antibodies (Table 1) were validated for AEC staining using known positive and negative control samples and stained through the Translational Pathology Shared Resource (TPSR) at Vanderbilt University Medical Center (https://www.vumc.org/translational-pathology-shared-resource). Between rounds of staining, imaging was performed using a Leica SCN400 slide scanner. In addition, chromogen and antibody stripping were also performed at the Digital Histology Shared Resource (DHSR) at Vanderbilt University Medical Center (https://www.vumc.org/dhsr).

**Table 1.**
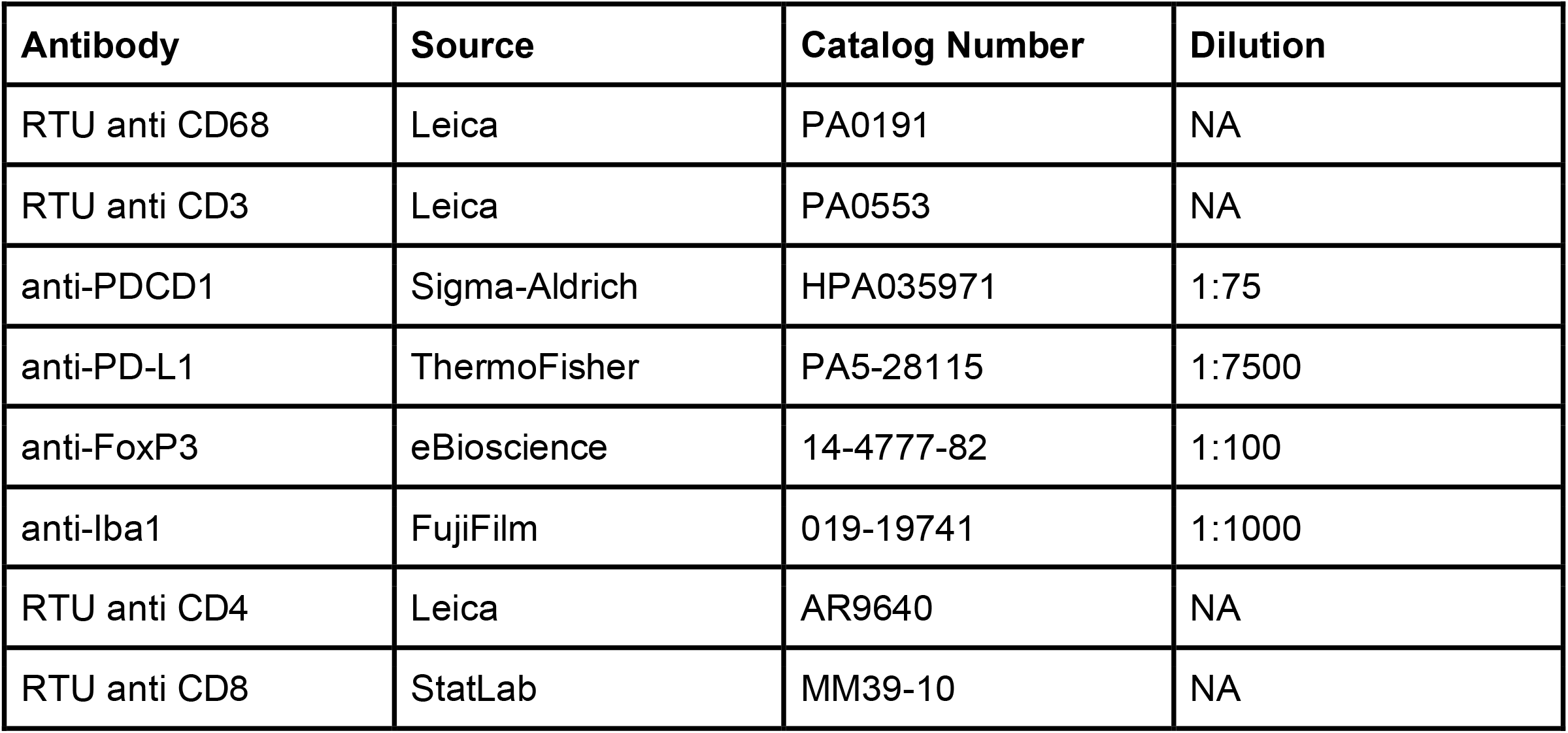

| Antibody | Source | Catalog Number | Dilution |
| --- | --- | --- | --- |
| RTU anti CD68 | Leica | PA0191 | NA |
| RTU anti CD3 | Leica | PA0553 | NA |
| anti-PDCD1 | Sigma-Aldrich | HPA035971 | 1:75 |
| anti-PD-L1 | ThermoFisher | PA5-28115 | 1:7500 |
| anti-FoxP3 | eBioscience | 14-4777-82 | 1:100 |
| anti-Iba1 | FujiFilm | 019-19741 | 1:1000 |
| RTU anti CD4 | Leica | AR9640 | NA |
| RTU anti CD8 | StatLab | MM39-10 | NA |

Whole slide image data import was performed using open source python packages tifffile, czifile, and fcsparser. Whole slide images were processed using open source python packages including scikit-learn and scipy. Data visualization was performed using matplotlib, seaborn, and napari. Multiprocessing was performed using packages including joblib, itertools, and multiprocessing. Segmentation was performed using a custom trained Stardist machine learning model. Keypoints registration was adapted from code developed during the 2020 NIH Cancer Systems Biology Center / Physical Sciences - Oncology image analysis hackathon (8). The full CASSATT pipeline and demo dataset is available at https://github.com/ihrie-lab/CASSATT. Image structural similarity and normalized mutual information were calculated using the metrics in the python scikit-image package.

Techniques for collection and analysis of the external cyclic IHC dataset can be found in Thomas et., al 2021 (9).

## Results

To maximize data collection per experiment, CASSATT was designed to process images from slide scanning microscopes that capture the entire slide rather than specific regions of interest. It can also be modified to process tissue microarray (TMA) data simply by passing each TMA core image as a ‘slide’ to the pipeline. Images should be acquired at sufficient magnification to clearly distinguish single nuclei using a standard hematoxylin stain - typically, <1 micron/pixel. Scanned images from most standard slide scanning microscopes can be directly imported into the pipeline. Accepted file types include TIFF, BigTiff, SCN, SVS, and CZI.

Throughout the pipeline, some intermediate files are saved to disk. Doing so decreases system memory usage and also allows the user to verify the success or failure of specific pipeline steps by opening and inspecting intermediate outputs. We have also implemented a ‘development mode’ which when enabled will save or display to console many more intermediate outputs, allowing users to quickly tweak and update settings for their dataset. This iterative quality control process is crucial in order to successfully analyze datasets accurately. Users can also set a seed value for the stochastic dimensionality reduction tools and clustering tools, ensuring a reproducible result given identical input data.

To analyze single cell expression data from a cyclic IHC experiment, near pixel perfect registration across all of the rounds of staining is crucial. This pipeline accomplishes this goal in two steps. First, rounds of staining are registered at a lower resolution that captures the presence and shape of the full tissue section (**Figure 2A**). Registration at this level is more robust to large scale shifts in the tissue position within the imaged area, as well as loss of tissue due to repeated handling of slides required for cyclic staining. In addition, registration of lower resolution images is much less computationally taxing than attempting to register native resolution images. A Euclidean transformation matrix consisting of translation values in x and y dimension and rotation angle is calculated on the lower resolution image. It is then upscaled by multiplying the translation values by the scale factor and is applied to the native resolution image.

**Figure 2.**
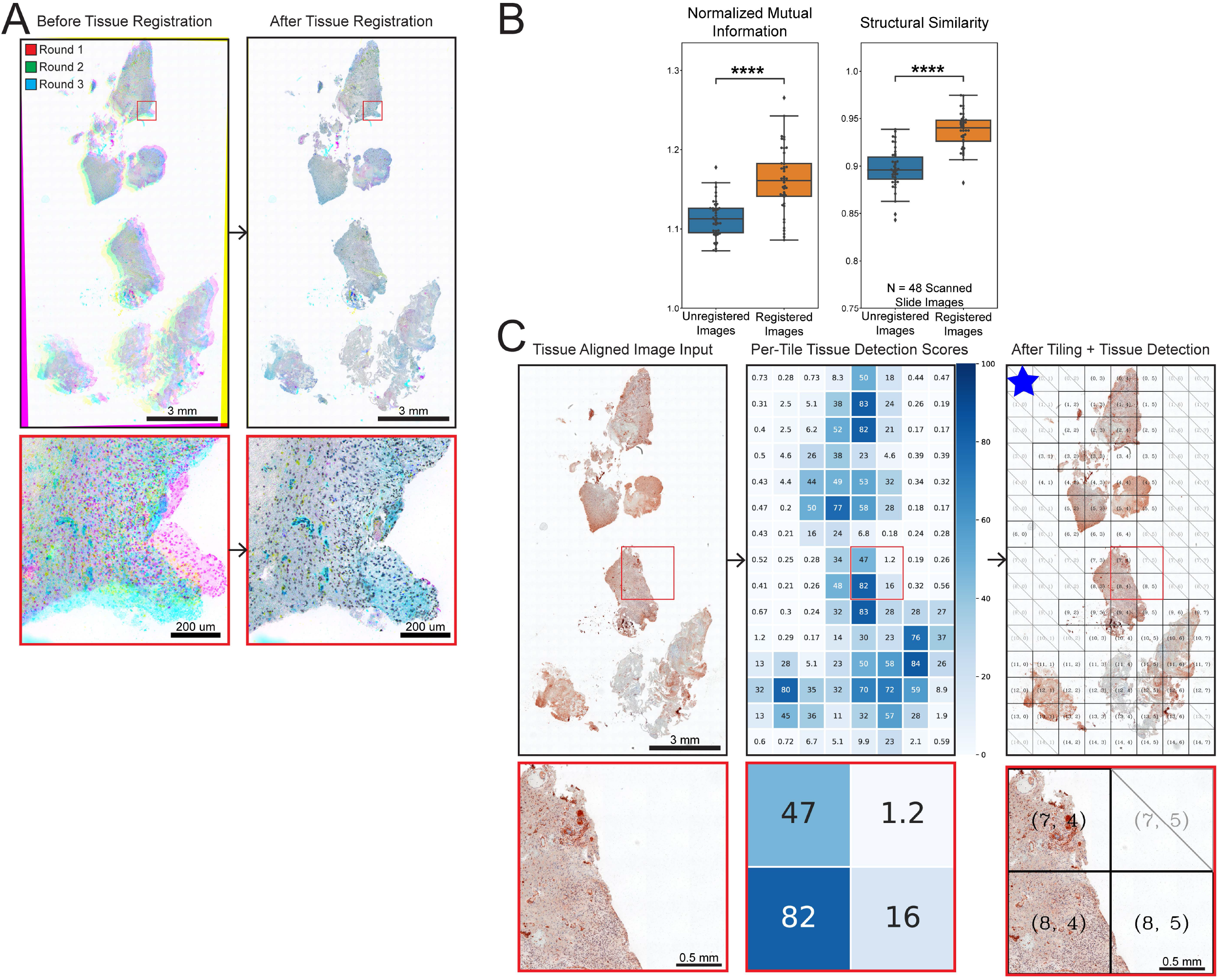
Tissue Registration and Tiling (A) Images from three rounds of imaging were flattened to grayscale and displayed together as the three channels of an RGB image to highlight round to round registration errors typical to cyclic IHC staining. Full slide view of tissue fragments and higher resolution view of a region of interest (red outline) are shown. (B) Normalized mutual information and structural similarity of pairs of images before and after tissue registration are plotted. Paired t tests with Bonferroni correction were performed (N = 48 scanned slides − 6 slides, 8 rounds. p = 2.19e-10 and p = 1.14e-9 respectively). (C) Post tissue registration image is broken into tiles and percent of tissue on each tile is scored. Only tiles where sufficient tissue is detected are saved for further processing. Blue star icon indicates a default CASSATT output saved to disk.

The reliability of tissue registration was tested by introducing random pixel translations in x and y dimension in addition to random small angle rotations to the test dataset images. Normalized mutual information and structural similarity of each image and its corresponding round zero image was recorded before and after tissue registration. Both metrics of image similarity showed significant improvement after tissue registration was performed (**Figure 2B**). We did not expect these metrics to reach perfect similarity because the rounds of images are not identical. Most importantly, the presence of a different biomarker stains for each round and tissue loss contribute to the differences between rounds. In addition, tissue registration aims simply to overlay like for like tissue regions and is not designed to achieve pixel perfect registration.

Following tissue registration, each image is divided into tiles of user-defined proportions. An ideal tile size may depend on several factors including the size and proportion of input scanned images, the computational resources available, and size of tissue features. Image tiling allows the subsequent analysis steps to run more quickly without over taxing the system. Tiles include a user-defined tile overlap parameter that helps ensure that when tiles are pixel registered and reassembled, any gaps on the perimeter of the tile are filled by the overlapping region. During the tiling process, a tissue detection algorithm is applied to each tile and only tiles where sufficient tissue is detected are saved for further processing (**Figure 2C**, middle). A low resolution tilemap with tile boundaries and row, column identifiers overlaid is saved as an output for each round’s image (**Figure 2C**, right). This tilemap can be used to visually identify specific tiles of interest and can also be used to quickly verify the success of tissue registration throughout the dataset. Additionally, a spreadsheet of tissue detection scores for each tile is exported.

Following tissue registration and tiling, each of the tiles where tissue was detected across all rounds of staining are collected for pixel level cell registration. To avoid registration errors caused by differences in marker expression between rounds, the hematoxylin and AEC stains are computationally unmixed using the separate_stains module from skimage, and only the hematoxylin stain is used for a keypoints detection and matching registration strategy. In each unmixed hematoxylin image, keypoints are detected and their coordinates and descriptors are saved. Each image is first registered to its corresponding round zero image. Registering all images in a series back to the original round image eliminates the danger of a ‘drift’ in registration where registration errors may be propagated down the line of registrations. To register two images, a matching algorithm is used to find keypoints with similar descriptors. This list of matches generally contains a large number of ‘correct’ matches and a smaller number of ‘incorrect’ matches that arise because of random chance. A transformation is computed that minimizes the distance between matching keypoints and maximizes the number of keypoint matches that produce this transformation (**Figure 3A**). The cell-level registration typically corrects for the 1-5 pixel misalignments that remain after the high level tissue registration step, and produces significantly more accurate registered images (**Figure 3B, C**).

**Figure 3.**
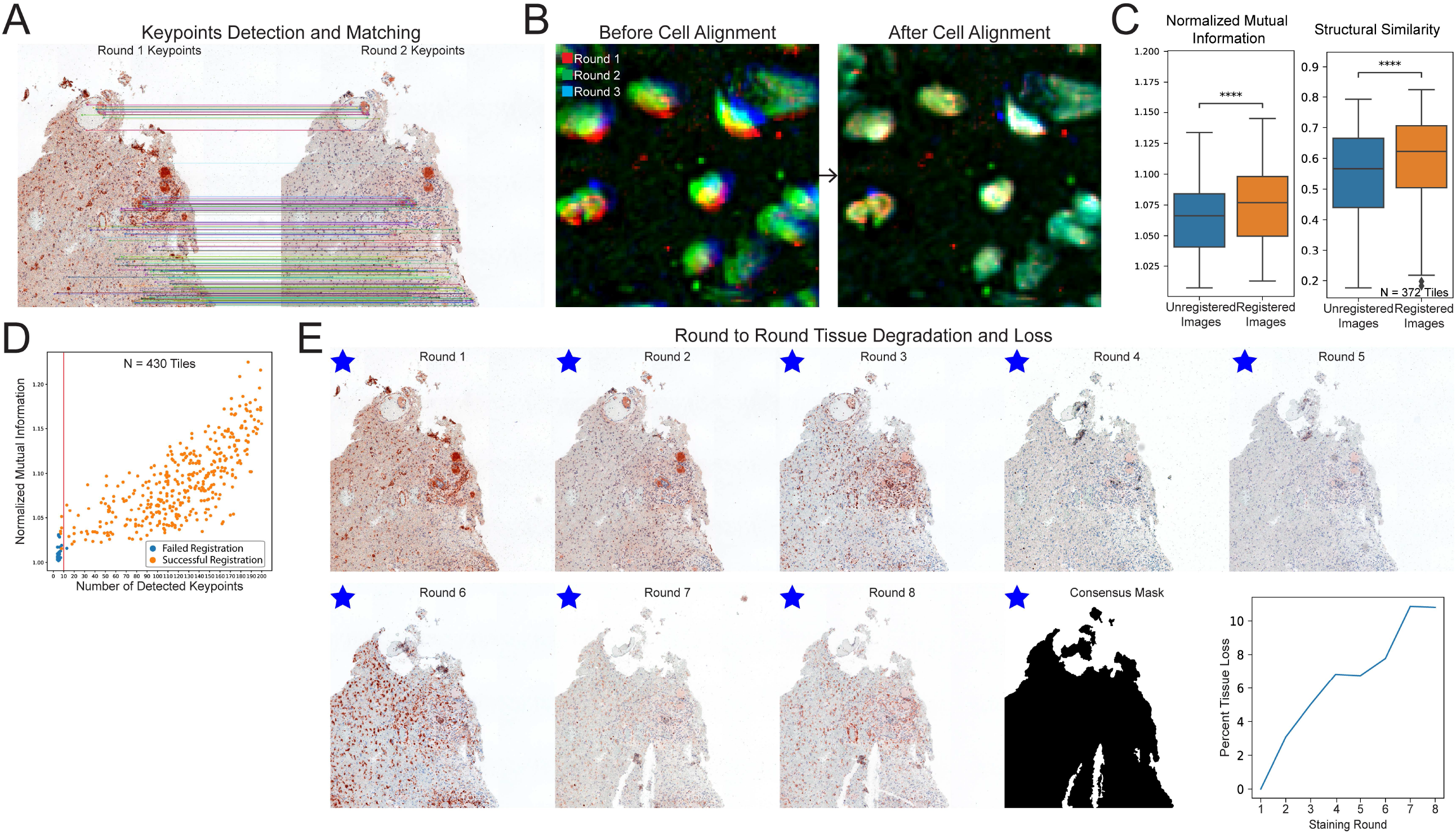
Cell Registration, Tissue Detection, and Masking (A) Successful keypoint matches are indicated by encircling the keypoint on each image, with a line drawn between the matching keypoints to demonstrate the registration. Many parallel matching keypoints indicate a high quality keypoints registration. (B) Unmixed hematoxylin images from three rounds of staining are overlaid as an RGB image to highlight round to round registration errors remaining after the tissue registration step. After cell alignment, these errors have been resolved. (C) Normalized mutual information and structural similarity of pairs of images before and after tissue registration are plotted. Paired t tests with Bonferroni correction were performed (N = 372 tile images. p = 1.56e-24 and p = 1.61e-38 respectively). (D) Plot of detected keypoint matches versus normalized mutual information of registered images. Default minimum matches to assume successful registration set at 10. (N = 430 Tiles) Ground truth registration failure and success manually scored. (E) Tile image is shown for every round of staining, Tissue degradation and loss can be seen in the top region of the image in round 4 and the bottom of the image in round 7. This damage and loss have been accounted for in the consensus mask which is then applied to all images. Plot of percent tissue loss across successive rounds of staining is shown.

The registration algorithm mathematically requires a minimum of four matching points to produce a transformation, but due to the possibility of false matches, the greater the number of detected matches, the greater the likelihood of a successful registration. A threshold value for the minimum number of keypoint matches required can be set by the user to automatically detect registration failures and exclude them from the dataset (**Figure 3D**). This threshold can be set based on a breakpoint in the NMI scores before and after registration, or by manually checking log files for the number of matches found in failed registration rounds. In the rare instances where the registration to the round zero image fails, the algorithm automatically attempts a second registration to the successfully registered image of the round immediately previous, and again checks for a successful registration. In cases where major tissue loss or distortion has occurred in a tile through the course of the experiment, this second registration can succeed where the first fails (**Figure S1**). If both registration attempts fail, the tile is eliminated from the analysis. For this reason, it is important to check that the failure thresholds are correctly tuned to prevent failed registrations from entering the dataset and successful registrations from being discarded. In addition, by performing this registration at the tile level, it is much more capable of correctly registering cases where areas of the tissue gain different distortions without necessitating the use of any computational warping of the image.

One of the drawbacks to any cyclical staining and imaging protocol is the inevitable tissue loss and damage that occurs throughout the rounds of staining and imaging. It is important to exclude any tissue areas that are lost from the analysis to prevent missing data or false negative data. To accomplish this, tissue detection is performed on each registered image and a consensus mask is produced in which only pixels where tissue was detected in every image in the stack are included (**Figure 3E**). This mask is applied to all of the images in the stack, and this final image stack is exported for further analysis. Users can also set a percent threshold of tissue loss per tile that is acceptable for their analysis. This can be used to automatically eliminate tiles where an inordinate amount of tissue loss has made the remaining tissue less useful for analysis.

After full pixel registration is complete, single cell nuclear expression data is generated by first segmenting nuclei on the unmixed hematoxylin signal and measuring the AEC signal overlapping each nucleus (**Figure 4A**). Color deconvolution is performed using the skimage implementation of the Ruifrok method for separation of hematoxylin and AEC stains (10). Cytoplasmic signal is similarly measured by expanding each nuclear mask by a user-defined number of pixels and measuring the overlapping AEC signal (**Figure 4B**). Segmentation is performed only on the first image in the stack and the outlines are applied to the full stack to avoid segmentation differences that can occur due to small changes in hematoxylin staining throughout the experiment. By default, nuclear segmentation is performed using a custom trained Stardist machine learning model for segmenting 2D hematoxylin images. Users can substitute their own trained Stardist model, or use any segmentation strategy that they prefer as long as it produces a nuclear segmentation mask that can be imported into python. The single cell expression data, paired with the positional coordinates for each detected cell is exported. (**Figure 4C**) This output can be analyzed in most single cell data analysis software or platforms. CASSATT enables two methods of cell population identification. First, a user can gate populations of interest in a cytometry program of their choice and read those populations back into CASSATT (**Figure 4D**). Alternately, CASSATT can take the expression data and perform dimensionality reduction and clustering to produce cell populations in an unsupervised fashion (**Figure 4E**). These methods will likely produce divergent cell populations, thus one or both options may prove useful depending on the dataset. These cell populations serve as the starting point for neighborhood analysis performed by CASSATT.

**Figure 4.**
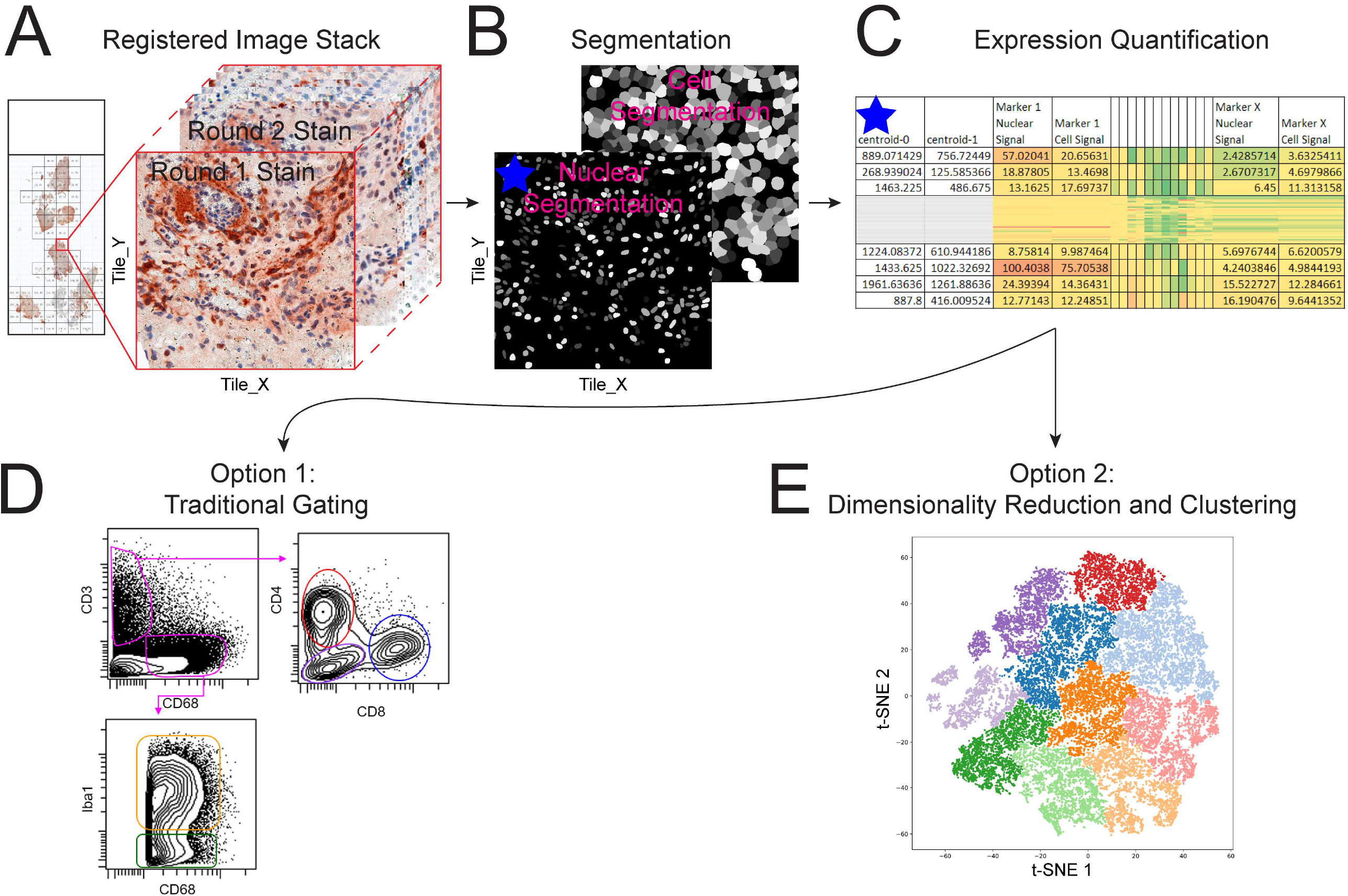
Segmentation and Gating (A) An image stack is assembled for each tile consisting of the corresponding image from each staining round. (B) Nuclear segmentation is performed using a custom Stardist segmentation model, followed by an estimated cell segmentation by pixel expansion. (C) Marker expression is tabulated and exported. Cell populations are identified by either traditional gating schemes (D) or by utilizing unsupervised dimensionality reduction and clustering techniques (E)

In order to analyze the spatial relationships between groups of cell types and their arrangement into specific niches, the workflow first must define how the neighbor cells of any specific cell are chosen. CASSATT includes a choice of three different methods for neighbor cell determination, each with certain advantages and disadvantages. In all cases, the central cell whose neighbors are being identified is referred to as the ‘index cell’. The first method makes use of the voronoi diagram that is produced by using the coordinates of every cell in a field as seed points and assigning every pixel to its nearest seed point. Thus, a cell’s voronoi neighbors are defined as the collection of cells that are the nearest seed point in the voronoi diagram in every direction from the index cell (**Figure 5A, B**) (4). In the second option, shell neighbors are defined as any cell whose coordinates fall within a distance set by the user (**Figure 5C**). Third, k-nearest neighbors are defined as the user-defined number of neighbors that are nearest in distance to any index cell (**Figure 5D**). The advantage and disadvantage of voronoi neighbors is that it returns a similar number of neighbor cells across cell densities. When cell density greatly decreases or at edges of tissues, voronoi neighbors will return neighboring cells that span gaps in the tissue or that may be too far away to be relevant. Shell neighbors will never include extremely distant cells but will return highly variable numbers of neighbors depending on the local cell density of index cells. Similar to voronoi neighbors, k-nearest neighbors are robust to density changes, however k-nearest neighbors can also return distant neighbors and can entirely exclude nearby cells in one direction from the index cell if there is a cluster of nearby cells in the other. In addition to these caveats for neighbor cell detection, it is also important to consider that all methods are only detecting neighbor cells in the 2D plane of the section, and this is necessarily a limited view of the niche in which a cell exists in 3D space.

**Figure 5.**
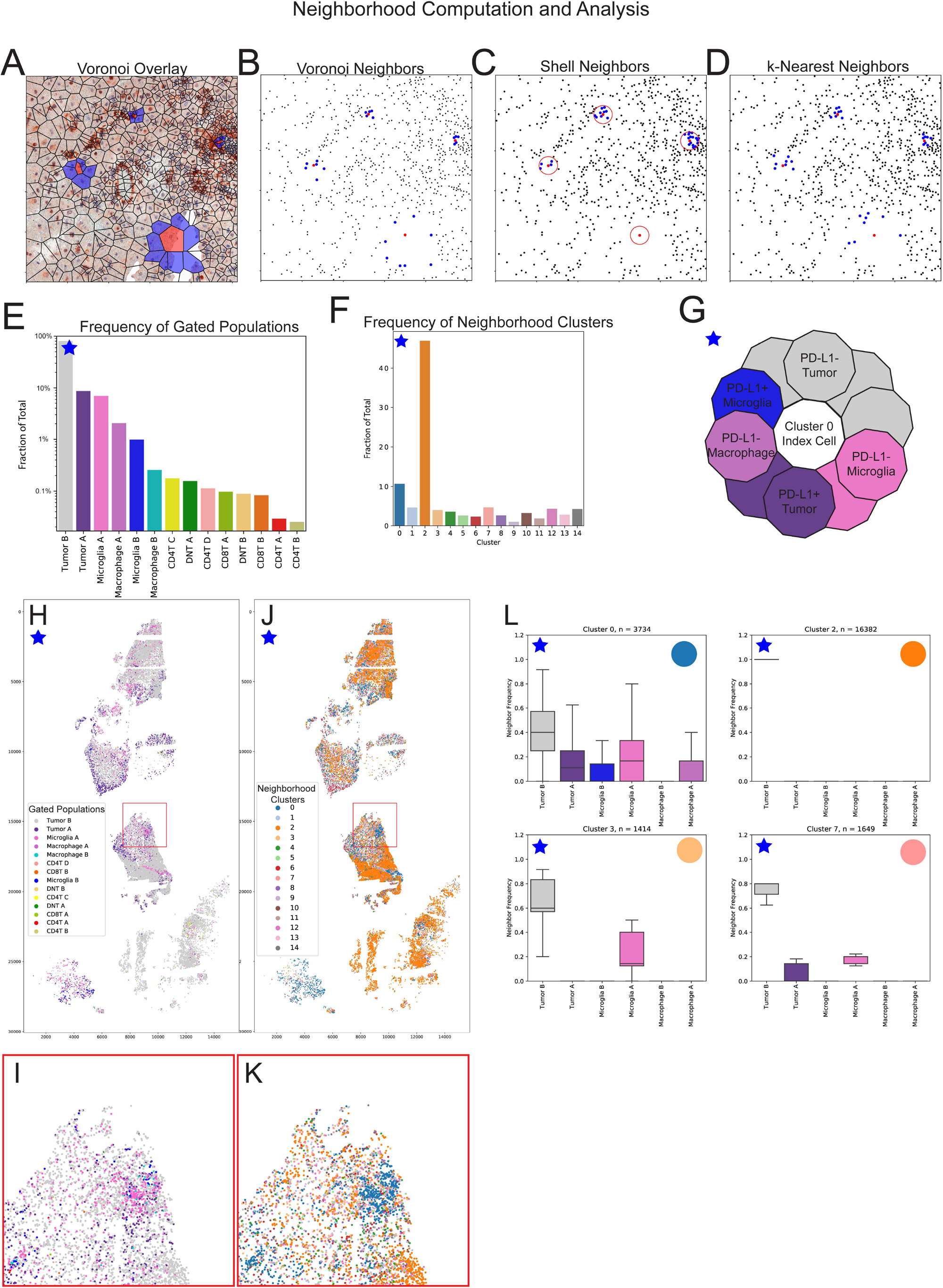
Neighborhood Computation and Analysis (A) A sample voronoi outline plot is shown overlaid on a bright field image. (B-D) Three strategies (voronoi, shell, and k-nearest) for neighbor cell identification are depicted, with each index cell shown in red and corresponding neighbor cells in blue. (E) Frequency of gated population identities across the full slide. Log scale is used to make low abundance populations visible. (F) Frequency of each neighborhood cluster across the sample is displayed as a percentage. (G) Approximation of neighborhood cells for an index cell of cluster 6 assuming ten neighbor cells. (H, I) Full slide of detections plotted and colored by their individual population identity and inset. (J, K) All neighbor cluster identities plotted from the test slide. (L) Frequency of each cell identity among the neighbor cells from each neighborhood cluster. Gated populations for which the 90th percentile neighborhood score was 0% for all neighborhood clusters are automatically dropped. For each plot, n = the raw number of index cells that compose the cluster.

To search for patterns in the neighborhood composition of cells, for each cell in the dataset we identify neighbor cells and, based on the cell subset identity of each cell’s neighbors, can calculate the relative frequency of each cell subset among the neighbor cells. These neighbor frequencies are the features that are used for dimensionality reduction and clustering resulting in clusters of cells with similar neighbor cell composition. The number of clusters found is set manually by the user. Additionally, a .csv file is produced containing the neighbor frequency of each cell subset for every detected cell. Importantly, the cell subset identity of the index cell is not taken into consideration for this analysis. The frequency of gated cell subsets and neighborhood clusters within the sample is often central to interpreting neighborhood analyses (**Figure 5E, F**).

The location and cell subset identity of all cells from a slide can be displayed to visually distinguish large scale patterns of subset localization (**Figure 5H, I**). Similarly, the cells belonging to each neighborhood cluster can be displayed (**Figure 5J, K, S2 A**). To better understand the neighbor cell makeup of each neighborhood cluster, a series of plots are produced. A hierarchical clustering diagram of each neighborhood cluster and the frequency of each of the cell populations shows which neighborhood clusters are most closely related (**Figure S2 B**). If there are clusters that are very closely related, a cut point can be set by the user on the hierarchical clustering tree to produce meta-clusters. To provide a high level visual representation of the neighbor cells that make up the neighborhood for any one neighbor-cluster, CASSATT produces a diagram that displays the assortment of neighbor cells that would result from the mean neighbor frequency of each gated population assuming ten neighbor cells (**Figure 5G, S3**). For each neighborhood cluster, a boxplot is generated that shows the distribution of neighbor frequency for each cell subset (**Figure 5L, S4**). A cluster with narrow box plot ranges indicates that the niches within that cluster have a high degree of purity and consistency in terms of the neighbor cell frequencies. On the other hand, wide ranges of neighbor frequencies with a cluster indicate that there is a high degree of heterogeneity in neighbor cell frequency with that cluster. For example, cells that fell in cluster 2 had uniform neighbor cells, consisting 100% of Tumor_B cells. Clusters 3 and 7 are examples where a relatively small fraction of immune cells compared to tumor are present in the vicinity of the index cell, while cluster 0 neighborhoods exhibit a higher frequency of a variety of immune cells. Comparison of the plots in Figure 5H and 5J shows that cluster 2 cells are located in the regions of the slide where Tumor_B cells are the overwhelming majority. Cluster 0, on the other hand, appears to consist of half and half Tumor and Immune subset cells, and comparison of Figure 5I and 5K demonstrates that blue cluster 0 cells correspond to areas of high immune infiltration.

In addition to finding clusters of cellular neighborhoods within the dataset, CASSATT will also calculate the log odds ratios of interactions between gated populations. Log odds values represent the relative chance that cells of two populations will interact with each other given their frequency and distribution within the dataset. A log odds value greater than 1 indicates a greater than random chance interaction frequency, and conversely a log odds value less than −1 indicates the interaction frequency is less than what would be expected by random chance. Log odds are calculated for every population pairing, and the raw number of interactions the log odds value is based on is also reported (**Figure S5 A**). This output allows a user to systematically search for potentially biologically significant interactions between cell types within a sample given that they occur at rates not predicted by chance. Population pairings where no interactions were found are left blank. To determine if the log odds interaction patterns differ from region to region within a tissue, in addition to a slide-wide calculation, per-tile log odds interaction heatmaps are also produced (**Figure S5 B**). Finally, to determine if correlations exist such that for example, a high log odds value for one population interacting with another population would predict a high log odds value for another population interaction pairing, a correlation matrix can be produced for each interaction pairing versus every other interaction pairing (**Figure S5 C**). Positive correlation values mean that across the tiles, positive log odds of interaction A correlated with positive log odds of interaction B. Negative correlation values mean that across the tiles, positive log odds of interaction A correlated with negative log odds of interaction B. Given an interaction of interest, this output allows a user to determine if that interaction correlates with any other interactions in the dataset.

CASSATT was tested on an externally published 18 round AEC multiplex IHC dataset to confirm consistency of results with prior analysis techniques. This dataset includes 208 total tissue microarray cores, representing 55 SCLC cases and 21 control cores including human tonsil tissue. Given the immune focused staining panel used, a tonsil tissue core was chosen to visually illustrate CASSATT outputs (**Figure 6A**). A selection of stains combined with the nuclear segmentation mask can be used to visually identify regions of the test core in which T-cells (CD3, CD8), B-cells (CD20), and dendritic cells (CD11c) are present (**Figure 6B**). Major immune populations were expert gated from the AEC signal quantified on each cell in the dataset (**Figure 6C**). These cell types were mapped back onto their original coordinates across the entire slide, and our specific test core (**Figure 6D**). Finally, the gated cell identity of cells was summarized per core and the tissue cell density was calculated. This density value was compared for each cell type in each core to those identified by Thomas et al (9). A Pearson correlation coefficient of 0.949 demonstrated a high degree of correlation, further supporting CASSATT’s utility and agreement with prior quantification techniques.

**Figure 6.**
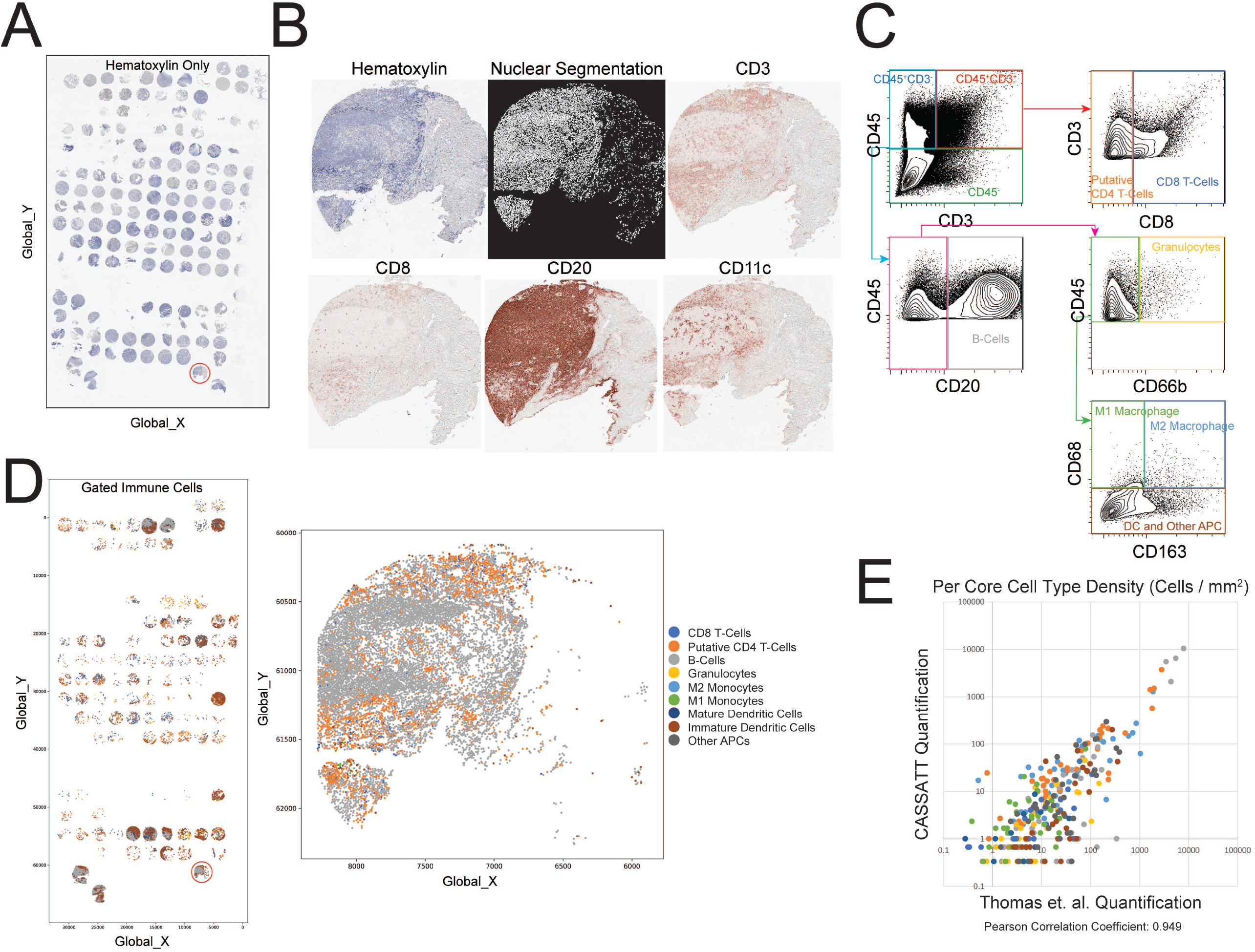
CASSATT Validation on Secondary TMA Dataset of Small Cell Lung Cancer (SCLC) Tumor Samples (A) Overview of hematoxylin stained TMA slide with sample core encircled. (B) Nuclear segmentation mask and subset of sample stains. (C) Biaxial gating scheme for major immune populations. (D) Cells colored by gated immune population identity across slide, and specific sample core. (E) Comparison of per core cell type density (cells/ mm^2^) calculated by CASSATT and Thomas et. al. (9)

## Discussion

As quantitative single-cell biology advances, an increasing number of parameters can be used to identify specific cell subtypes of interest. Discovery datasets may include 35, 50, or more parameters, including both antibody-dependent (immunostaining) and native features (e.g., nuclear size and eccentricity). To adapt the findings from discovery research to lower-dimensional tests on large numbers of samples, key parameters are often identified and then confirmed using standardized immunostaining approaches with colorimetric or fluorescent detection. However, even within lower-dimensional studies, more than one antibody-based label is often needed to find cells and patterns of interest. Within the immune system, which is studied in this test dataset, it is increasingly common for a combination of multiple proteins to be required to distinguish functional subtypes. In the case examined here, beyond identifying lymphoid and myeloid cells using 4 markers, expression of PD-1 and its ligand PD-L1 is also needed to identify populations of cells likely to respond to immune checkpoint inhibitor therapies. Similarly, in stem and progenitor cell studies, a combination of 3 to 7 or more proteins may be required to finely distinguish the specific population of interest (11-15). For this number of parameters, cyclic immunohistochemistry can be a rapid and largely automated approach that can be performed on a large number of samples in parallel. CASSATT is designed to provide similarly streamlined analysis of the data resulting from these stains.

Depending on the dataset and processing step, CASSATT’s speed can be rate limited by a workstation’s memory or CPU access. Across the workflow, three steps represent the lion’s share of processing time - tissue registration, cell registration, and neighborhood cell computation. For steps where memory is not rate limiting such as per-tile pixel level registration and per-tile neighbor cell calculation, CASSATT utilizes parallel processing to distribute jobs across 90% of available cores. In our test dataset on a 32 core workstation, this parallelization unlocked approximately 25X faster registration of all tiles. On the other hand, tissue registration on full slides is highly memory limited so CASSATT defaults to processing one job at a time. However, in the case of TMA cores each being run as a ‘slide’ CASSATT allows the user to manually increase the number of parallel jobs up to the workstation’s core count.

CASSATT’s run time is highly dependent on the input dataset size. CASSATT’s run time was compared between two slides in the dataset representing ‘small’ and ‘large’ ends of the size spectrum (12 GB and 40 GB total) (Table 2). Unsurprisingly, CASSATT’s run time was significantly longer for the larger dataset, however when component steps are examined it is clear that the image processing steps of registration and segmentation contributed the lion’s share of the increased processing time while the neighborhood analysis did not take proportionally longer for the large dataset.

**Table 2.**

| Workflow Step | 12 GB Dataset | 40 GB Dataset |
| --- | --- | --- |
| Tissue Registration | 4 min | 20 min |
| Tiling | 1 min (73 Tiles) | 3.5 min (173 Tiles) |
| Cell Registration | 101 min | 25 min |
| Segmentation | 0.5 min | 1.75 min |
| Neighborhood Analysis | 23 min | 30 min |

The availability of three different neighbor identification approaches allows researchers to tailor the analysis to a specific research question. For example, studies of interactions that require tight cell-cell contact, such as Notch ligand signaling within a stem cell niche, or studies of proximity to a defined anatomical region, may be better suited to a shell method for defining close cellular neighbors. Alternatively, analyses of tissues with highly variable nuclear density and cellular shapes, including some malignant solid tumors and stratified epithelia, may be more amenable to a voronoi neighbor identification. Finally, k-nearest neighbors can be especially well suited to tissues with variable nuclear density where users want to expand or contract the range of neighbor cells used for statistical analyses.

As multiplexed spatial imaging and sequencing approaches proliferate, it will be useful to extend this pipeline approach to include additional imaging modalities such as dual-chromogen immunohistochemical staining or cyclic immunofluorescence. Each of these data types, while similar to the AEC chromogen detection used here, will require platform-specific background correction and optimization. More broadly, adaptation of segmentation and neighbor analyses to three-dimensional datasets, either from lightsheet imaging or three-dimensional imaging mass cytometry, will enable additional uses of this workflow in additional data types.

## Acknowledgements

The authors thank Dr. Joe Roland of the Digital Histology Shared Resource at Vanderbilt University Medical Center (www.mc.vanderbilt.edu/dhsr) for beneficial discussions and advice on the processing strategy for tiling and registration, and CSBC/PS-ON image analysis hackathon organizers and participants for inspiration and advice on registration and visualization techniques. The authors thank the members of the Ihrie and Irish labs for helpful discussions, and Giovanney Gonzalez (OHSU) for expert technical support related to immunohistochemical staining. Research in the Ihrie lab is supported by the National Institutes of Health (R01NS118580 and a supplement to U54CA217450 to R.A.I. and J.M.I.), the Ben & Catherine Ivy Foundation (R.A.I.), and a gift from the Michael David Greene Brain Cancer Fund at the Vanderbilt–Ingram Cancer Center (R.A.I. and J.M.I.). Research in the Irish lab is supported by the National Institutes of Health (R01CA226833 and U54CA217450) and the Human Immunology Discovery Initiative of the Vanderbilt Center for Immunobiology.

Work in the Lovly lab is supported by: a Vanderbilt Ingram Cancer Center Young Ambassadors Award, a Lung Cancer Foundation of America and International Association for the Study of Lung Cancer Lori Monroe Scholarship, and by the National Institutes of Health (NIH) (grant numbers U54CA217450-01, U01CA224276-01, P30-CA086485, and UG1CA233259). Dr. Thomas was supported by grant S21MD000104 and the Ann Melly Summer Scholarship in Oncology. Dr. Bartkowiak was supported by the National Cancer Institute (K00CA212447)

Dr. Coussens was supported by the National Institutes of Health grant P30-CA0695533; Stand Up to Cancer – Lustgarten Foundation Pancreatic Cancer Convergence Dream Team Translational Research Grant SU2C-AACR-DT14-14; OHSU Brenden-Colson Center for Pancreatic Health.

The Translational Pathology Shared Resource is supported by NCI/NIH Cancer Center Support Grant 5P30 CA68485-19, and Shared Instrumentation Grant S10 OD023475-01A1.

## Author Contributions

A.A.B, J.M.I. and R.A.I. conceived of the project. A.A.B. generated code and constructed the pipeline. R.K. reviewed and tested the workflow and prepared code as a pip-installable package. T.B. selected antigens for cyclic immunostaining analysis and performed expert gating to identify cell subpopulations of interest in both datasets. P.L.T., S.S., C.B.B., L.M.C. and C.M.L. provided the validation dataset and expert gating of cell subsets for comparison and validation. J.M.I. and R.A.I. provided financial support and project direction. A.A.B. and R.A.I. generated the initial draft, which was reviewed, edited, and approved by all authors.

## Figure Legends

Figure S1

Cell Registration on Increasingly Poor Tissue Integrity

Successive registration attempts for a tile that exhibited major tissue degradation and damage through the rounds of staining. Rounds 2 through 6 successfully registered to the Round 1 image as automatically detected by the n_matches value falling above the defined threshold of 10. Rounds 7 and 8 failed to successfully register with n_matches = 8 and 6 respectively. The follow-up registration attempts to the most recent successfully registered image were successful (n_matches = 37 and 23 respectively) Green boxes indicate successful registrations passed to downstream analysis.

Figure S2

Neighborhood cluster characterization and localization

(A) Hierarchical clustering of neighborhood clusters based on neighbor frequency of each gated population identity. (B) Localization and relative density of each neighborhood cluster is shown by highlighting cells belonging to individual clusters over the depiction of all analyzed cells from the slide.

Figure S3

Cartoon neighborhood depictions

A representative set of neighbor cells for each neighborhood cluster is shown. Cells are organized based on their frequency starting with the top most decagon and moving clockwise. The number of neighbor cells of each cell type is calculated based on the median neighbor frequency of each cell type within the cluster.

Figure S4

Neighborhood purity plots

Box and whisker plots for each cell type for each neighborhood cluster show the degree of homogeneity or heterogeneity of neighbor frequency within each cluster, as well as the neighbor cell types that characterize each cluster.

Figure S5

Log Odds of Interactions

(A) Log odds of interactions between each pair of cell types is depicted on a color scale. Numbers within each box indicate the raw number of interactions between the two cell types across the dataset. Cell type pairings where no interactions were found in the dataset are left gray and blank. (B) Log odds plots are produced in the same fashion but only for cells within a specific tile. (C) Given the per tile log odds of interactions, a correlation matrix shows the likelihood of logs odds of one interaction to influence the log odds of another interaction for each cell type-cell type interaction. Correlations of interactions for a selection of cell type pairs is shown here.

