## Supplementary figures and images for "Alignment, Segmentation and Neighborhood Analysis in Cyclic Immunohistochemistry Data Using CASSATT"

### Supplemental Figures

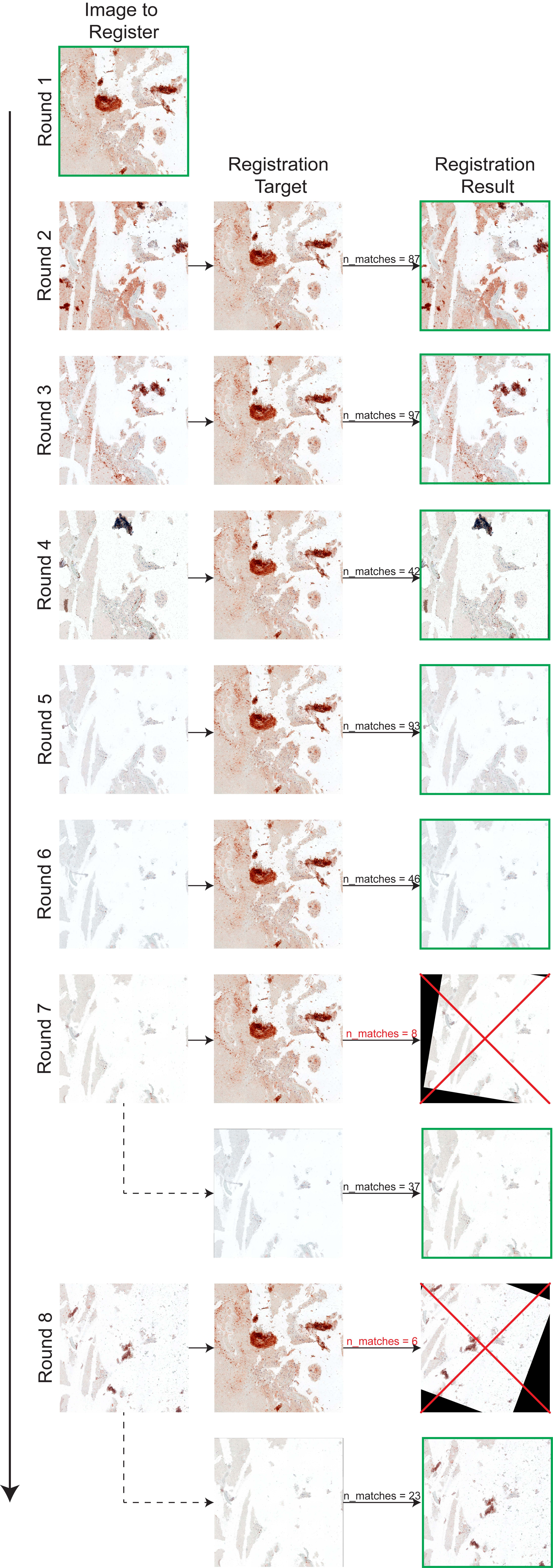

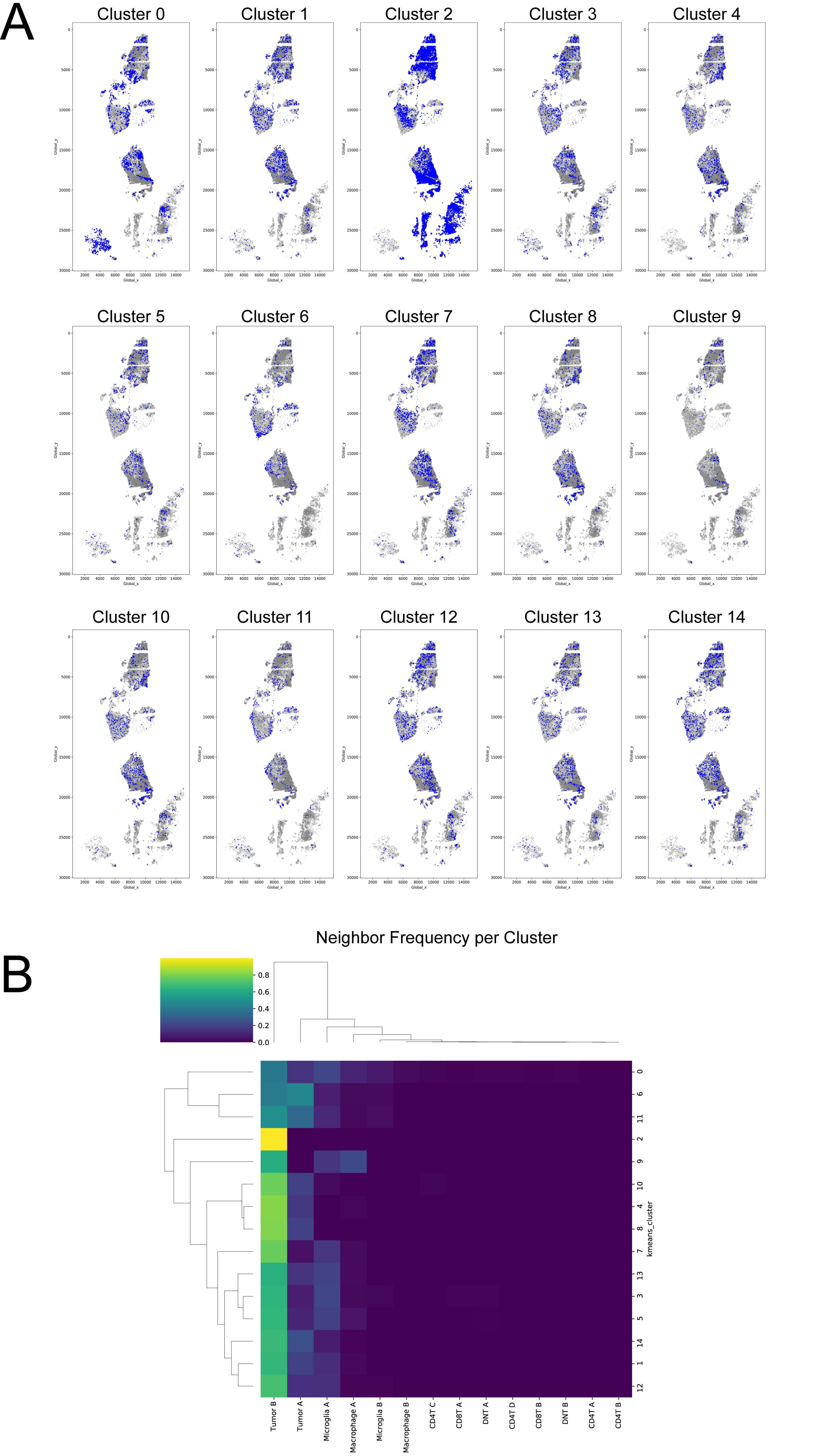

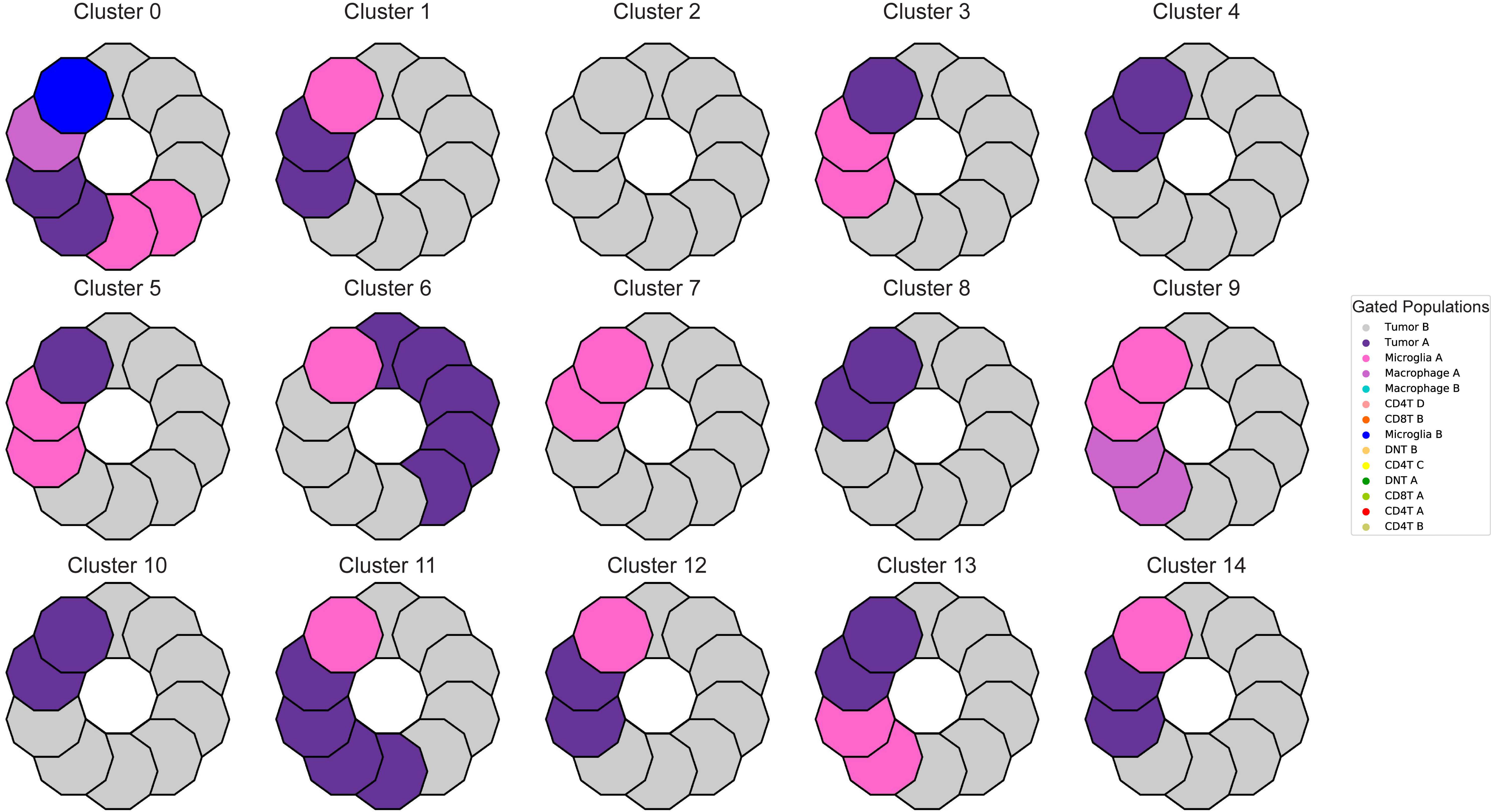

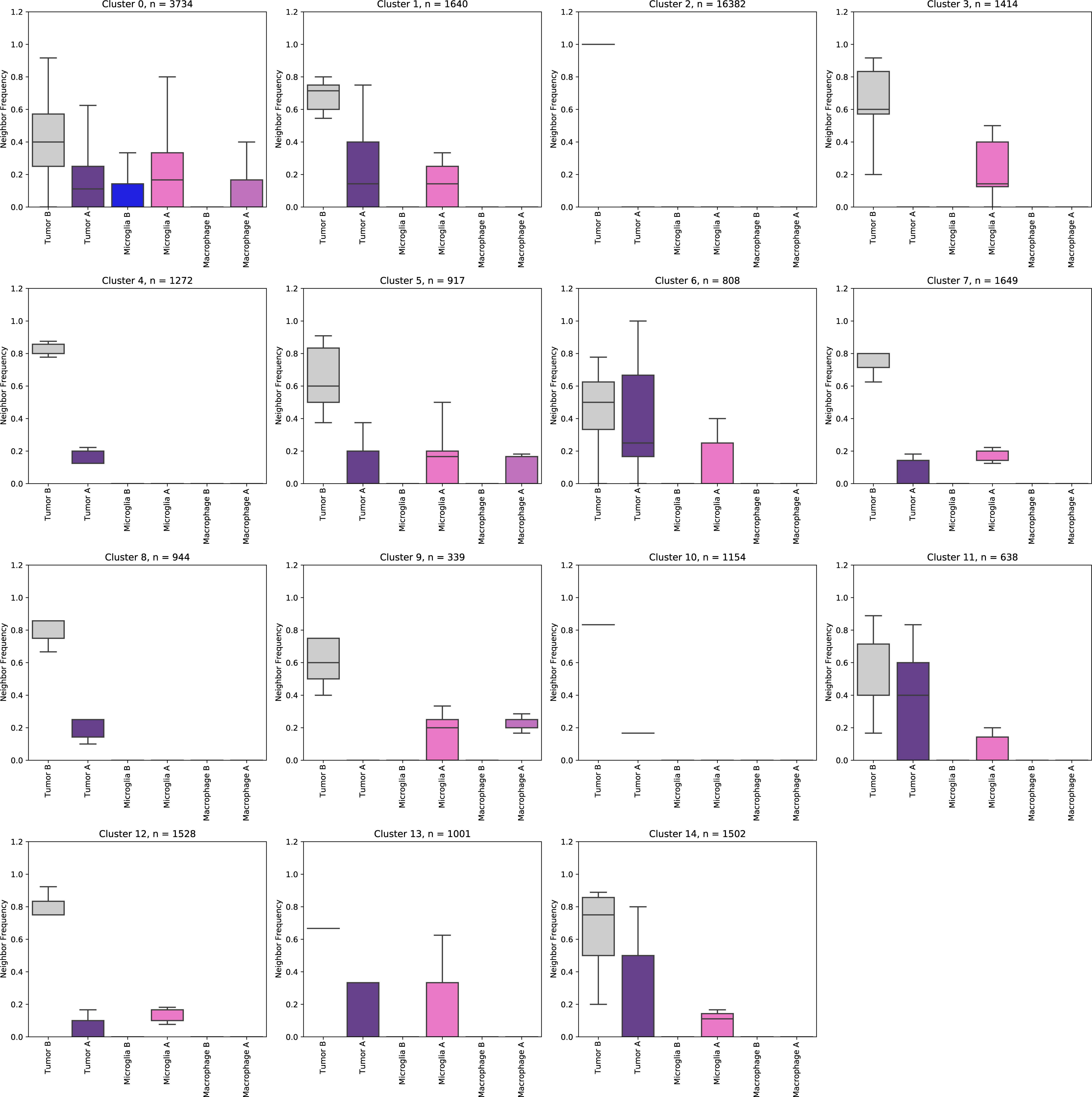

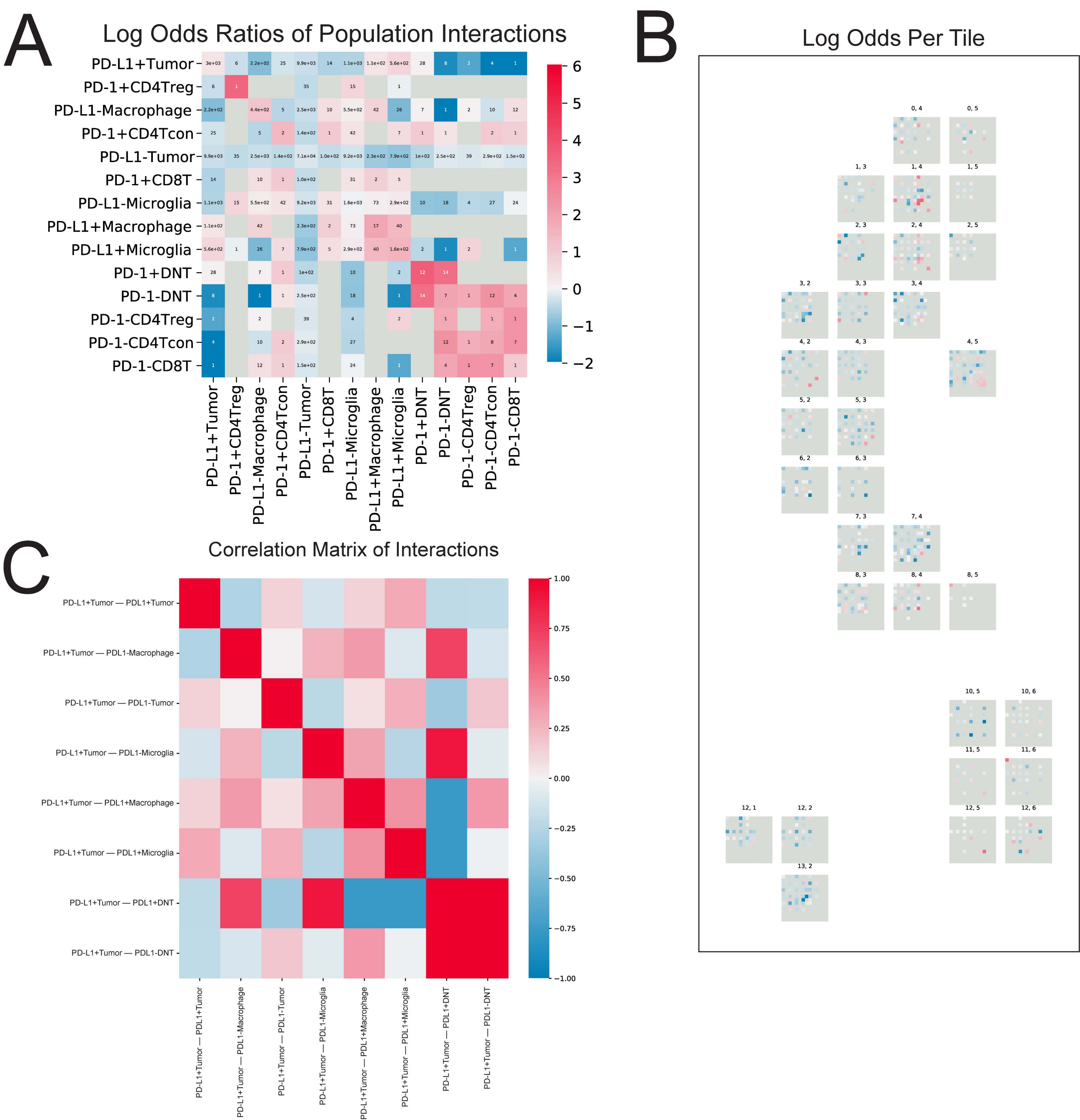
